# Combining Gamma Neuromodulation and Robotic Rehabilitation Restores Parvalbimin-mediated Gamma Function and Boosts Motor Recovery in Stroke Mice

**DOI:** 10.1101/2024.08.15.608060

**Authors:** Livia Vignozzi, Francesca Macchi, Elena Montagni, Maria Pasquini, Alessandra Martello, Antea Minetti, Éléa Coulomb, Anna Letizia Allegra Mascaro, Silvestro Micera, Matteo Caleo, Cristina Spalletti

## Abstract

Stroke is a leading cause of long-term disability, often characterized by compromised motor function. Gamma band is known to be related to Parvalbumin interneurons (PV-IN) synchronous discharge and it has been found to be affected after stroke in humans and animals. Both Gamma band and PV-IN also play a key role in motor function, thus representing a promising target for post-stroke neurorehabilitation. Non-Invasive neuromodulatory approaches are considered a safe intervention and can be used for this purpose. This study presents a novel, clinically relevant, non-invasive and well-tolerated sub-acute treatment combining robotic rehabilitation with advanced neuromodulation techniques, validated in a mouse model of ischemic injury. In the sub-acute phase after stroke, we scored profound deficits in motor-related Gamma band regulation on the perilesional cortex. Accordingly, both at the perilesional and at the whole-cortex levels, the damage results in impaired PV-IN activity, with reduced firing rate and increased functional connectivity levels. Therefore, we tested the therapeutic potential of coupling robotic rehabilitation with optogenetic PV-driven Gamma band stimulation in a subacute post-stroke phase during motor training to reinforce the efficacy of the treatment. Frequency-specific movement-related Gamma band stimulation, when combined with physical training, significantly improved forelimb motor function. More importantly, by pairing robotic rehabilitation with a clinical-like non-invasive 40 Hz transcranial Alternating Current Stimulation, we achieved similar motor improvements mediated by the effective restoring of movement-related Gamma band power and increased PV-IN connections in premotor cortex. Our research introduces a new understanding of the role of parvalbumin-interneurons in post-stroke impairment and recovery. These results highlight the synergistic potential of combining perilesional Gamma band stimulation with robotic rehabilitation as a promising and realistic therapeutic approach for stroke patients.

**Summary:** Stroke-induced motor deficits are accompanied by alteration of Gamma modulation and PV-interneurons activity and restored by a combination of non-invasive Gamma stimulation and robotic therapy.

## INTRODUCTION

Stroke-induced functional deficits in the motor cortex represent one of the main causes of disability worldwide. Recently, post-stroke physical therapy has been enriched by novel rehabilitative strategies, such as the combination of robotic tools with neuromodulatory techniques.(1–3) Robotic therapy is a promising tool for collecting highly precise kinetic and kinematic data about motor performance and providing customizable exercises. In addition, it is now widely accepted that physical therapy is crucial for stroke rehabilitation but should be coupled with neuromodulatory strategies to maximize its effectiveness. These strategies aim to boost plasticity and “prime” perilesional spared tissue, enhancing its susceptibility to “activity-dependent plasticity”. These strategies can involve plasticizing drugs (i.e. Fluoxetine) or Non-Invasive Brain Stimulation (NIBS) techniques that enable precise and controlled modulation of cortical activity.(4,5) The optimal interplay between a plastic environment and proper physical exercises is essential for motor recovery.(6)

Over the past decades, there has been a growing understanding of the importance of oscillatory neural activity in motor function in both healthy and injured subjects. Specifically, synchronized oscillations in the higher-Gamma frequency range (60–90 Hz), leading to an increase in Gamma power, coincide with movement initiation and execution (referred to as movement-related Gamma synchronization) and reflect the initial activation of primary motor neurons involved in movement.(7,8) Studies in humans have demonstrated that driving higher-Gamma oscillations enhances motor performance,(9) demonstrating a causal role of these neural rhythms in plasticity and motor control.(10,11) Conversely, lower gamma band oscillations (30–60 Hz) are implicated in strategies for controlling stronger muscle force production.(12)

In humans, an ischemic attack in the middle cerebral artery territory induces acute alterations of the neural rhythmic activity at rest in both affected and unaffected hemispheres.(13) Notably, an asymmetric enlargement of delta (2–3.5 Hz) and theta (4–7.5 Hz) band powers in the affected hemisphere accompanies ischemic events, while reduced gamma activity correlates with compromised hand functionality. Consistently, stroke survivors with incomplete upper-limb motor recovery and persistent deficits showed significantly lower Gamma-band corticomuscular coherence when performing a reach task using shoulder/elbow muscles.(14) This decreased coherence could reflect poor brain-muscle communication or poor integration of the signals from the two sources during motor actions. Poor EEG-EMG coherence could reflect an underlying mechanism contributing to impaired reaching performance in stroke patients.(14) Conversely, increased Gamma power in the affected hemisphere was associated with better recovery prospects.(13) Recent findings in animal models and human studies suggest a direct link between oscillatory activity in the Gamma band and the excitatory/inhibitory balance within reciprocally connected networks of GABAergic interneurons and pyramidal cells within the primary motor cortex.(15–17) Indeed, Nowak et al.(18) demonstrated that driving Gamma frequency oscillation via NIBS techniques is feasible and produces significant effects on motor function and GABA-mediated inhibition in healthy subjects.

In this context, specific firing characteristics of a well-defined subgroup of interneurons, the fast-spiking Parvalbumin-positive (PV) GABAergic interneurons, demonstrated a causal role in generating and modulating Gamma rhythms.(16,19,20) Recently, in rodent models for stroke, Parvalbumin activity has been related to post-stroke dysfunctions and to motor recovery.(21) Accordingly, fast spiking interneurons optogenetic modulation has been used to improve motor recovery in animal models.(22,23) Importantly, Wang and colleagues demonstrated that perilesional optogenetic 40 Hz stimulation on inhibitory interneurons in acute phase after stroke increases neuronal survival and plasticity(22,24). This body of evidence may provide a good foundation for the development of new therapeutic strategies involving the modulation of brain rhythms in association with physical therapy. Despite these advancements, a coherent framework integrating and adapting these novel paradigms into human post-rehabilitation strategies remains elusive. Optogenetic approaches and their very restricted timing have to face the clinical reality and must be properly translated for human application. This gap is also increased by the high variability among lesion volumes, infarct location and clinical approaches.(25) Moreover, the precise neurophysiological mechanisms underlying the effects of functional damage and post-stroke therapies are still poorly defined. As a result, despite rehabilitation, a significant percentage of stroke patients display persistent impairments in activities of daily living.

This study explores longitudinal effects of the ischemic lesion on PV-interneurons and movement-related Gamma band activity in a mouse model of ischemic stroke. Moreover, we used this data to validate the translational therapeutic potential of inducing gamma frequency oscillations in the spared forelimb premotor cortex combined with robotic-guided motor rehabilitation. We employed optogenetics to induce PV-IN-guided Gamma rhythm in combination with daily exercise on a custom-made device designed for mouse forelimb training (26). We then confirmed the efficacy of the treatment using a clinically relevant NIBS approach combined with robotic rehabilitation. Our data provide novel crucial insights into the neural mechanisms driving recovery and contemporary offers a reliable support for the development of more effective post-stroke therapies.

## RESULTS

### The ischemic lesion induces motor-related dysfunction in Gamma band synchronization

We first assessed the impact of an ischemic lesion in Caudal Forelimb Area (CFA) on Gamma band regulation in the spared perilesional premotor cortex (Rostral Forelimb Area, RFA) during voluntary movement. Mice had either undergone a photothrombotic lesion or a sham surgery in the CFA (Fig. 1A). A representative image of the stroke lesion is shown in Fig. 1B (dashed yellow line). Motor function assessment was performed with the Gridwalk test before and after stroke as shown in Fig. S1A. Local field potential (LFP) recordings from the RFA were taken while mice were engaged in a voluntary retraction task on the M-Platform, a custom-made apparatus for functional evaluation and neurorehabilitation of the forelimb in mice (Fig. 1C and 1D). We quantified Gamma band power within specific time windows delineating a baseline (i.e. −1 to −1.5 s from movement onset), a pre-onset phase (PRE, from −500 ms to onset) and a post-onset phase (POST, from onset to +500 ms) as illustrated in Fig. 1E. In sham animals, we observed a significant increase in Gamma band power in the premotor cortex in the pre-onset phase across all the cortical layers (Fig. 1F, grey bar plots). Throughout movement execution (POST), Gamma power remained elevated in the superficial layers (channel 1 to 8 of the linear multiprobe), while Gamma modulation was minimally evident in deeper layers of the cortex (channel 9 to 16 of the linear multiprobe, Fig. 1G, grey bar plots). In contrast, stroke animals assessed 2 and 5 days post-lesion exhibited no Gamma band upregulation before movement onset (Fig. 1F, light blue and blue, respectively) and showed significantly lower Gamma power during movement compared to controls. Consistently, during movement execution, Gamma band power in stroke animals decreased across all cortical layers and resulted significantly different from controls in upper layers, where Gamma band regulation was more pronounced (Fig. 1G). Baseline Gamma power did not differ between sham and stroke mice (Sup1B).

**Fig. 1.**
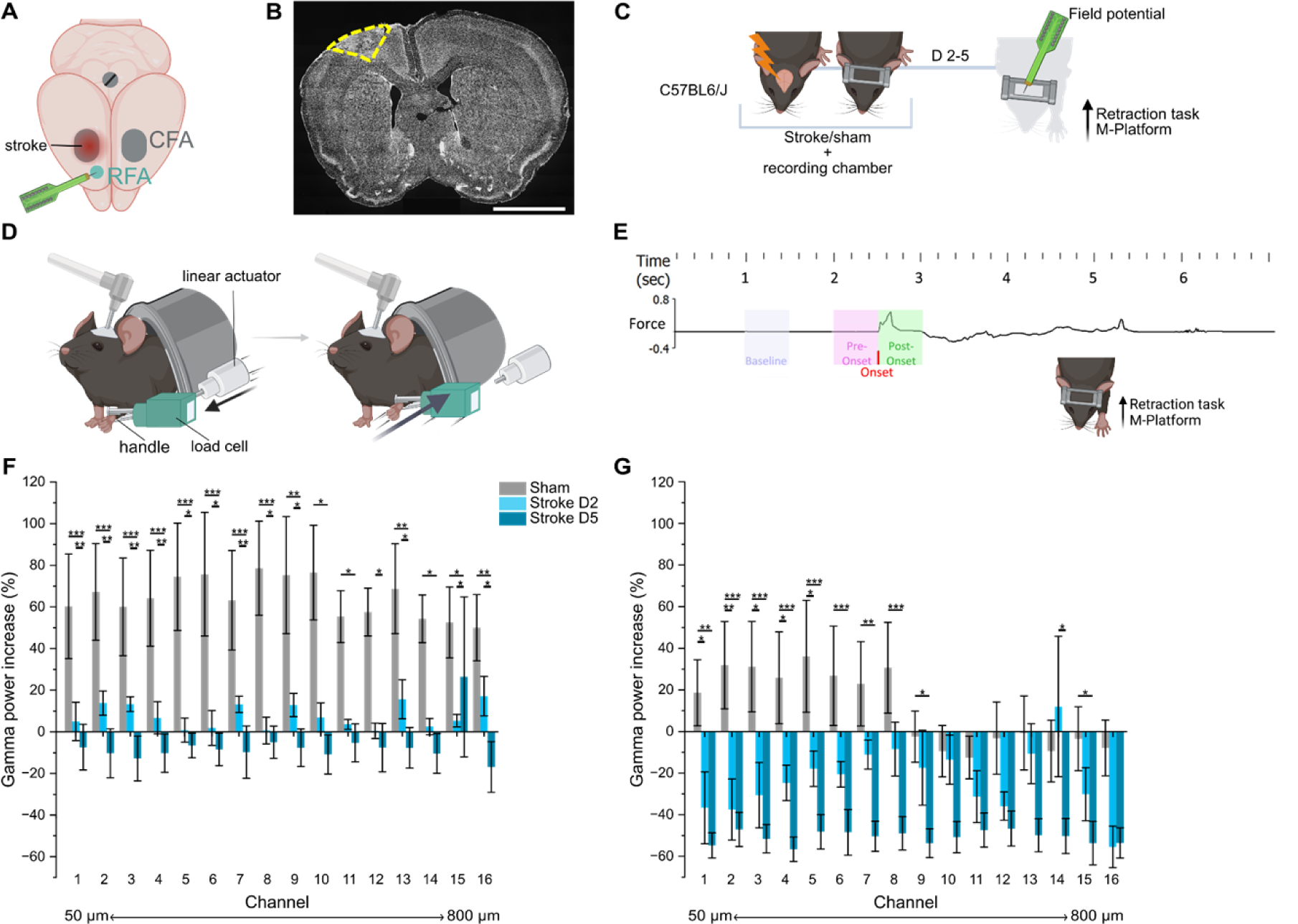
Impaired motor-related Gamma band regulation after stroke. Electrophysiological recordings during voluntary movement in the perilesional premotor cortex revealed a significant deficit in Gamma band modulation following ischemic stroke in the motor cortex. **A**, Schematic representation of the stroke lesion in the right CFA and the perilesional region (RFA) where the electrophysiological recordings have been performed. **B**, Representative coronal brain slice of a stroke animal labeled with Hoechst nuclear staining shows the lesion (contoured by the dashed yellow line). Scale bar 2 mm. **C**, Schematic representation of the experimental protocol. **D,** Schematic representation of the two different phases of the retraction task on the M-platform. Mice are head restrained with their wrist closed in a handle connected to the load cell for force detection. In the passive phase (left) the linear actuator pushes forward the handle and mouse forelimb. In the active phase (right) the mouse is trained to return the handle to the home position by overcoming a defined friction in order to receive a reward. **E,** Windows of analysis during the task in relation to movement onset. The black line represents an example of force trace throughout the task; the onset of the movement is highlighted in red; the purple window identifies the baseline period when no movement is detected; the pink window identifies the pre-onset phase (i.e movement preparation) and the green window the post-onset phase (i.e. movement). **F** and **G,** Quantification of Gamma band power across all the 16 channels spanning all the cortical layers (channel 1 ≃ 50 μm, channel 16 ≃ 800 μm) in pre-onset and post-onset windows, respectively. Healthy mice (*n =* 9) are shown in grey, animals recorded 2 days after stroke are shown in light blue (*n =* 6), while mice recorded 5 days after stroke are represented in dark blue (*n =* 8). The positive Gamma band modulation in pre-onset and in the higher layers in post-onset phase is abolished and even reversed in subacute stroke. Two-Way ANOVA followed by Tukey test, * *P* < 0.05, *** *P* < 0.001. Data are shown as mean ± SEM.

### Ischemic stroke profoundly changes resting state functional connectivity of PV-IN up to one month after injury

We reasoned that the impairment of Gamma band regulation could be associated with local and widespread changes in the functionality of PV-interneurons. To test this hypothesis, we performed longitudinal wide-field calcium imaging in awake head-fixed mice expressing GCaMP7f in PV-interneurons. We recorded PV spontaneous activity one day before (PRE-STROKE) and at 2, 5, 8, 14, 21 and 28 days after stroke in the same mice. Spontaneous PV activity underwent a drastic reduction in the peri-infarct cortex, involving a large fraction of both hemispheres (Fig. 2A). We therefore hypothesized major alterations in PV-IN functional connectivity (FC) after the lesion. Thus, FC of the entire dorsal cortex was evaluated by computing Pearson’s correlation (Fisher’s z-transformed) between all paired cortical regions after hemodynamic correction (Fig. 2B). In healthy mice, resting state FC was homogeneous and symmetrical across hemispheres. Notably, anterior primary motor cortices displayed low correlation values, with the sole exception being the homotopic connectivity. In the sub-acute phase after stroke, marked alterations were evident, with opposite consequences on intra- and inter-hemispheric FC (Fig. 2C). Although far from pre-stroke condition, the average FC became more homogenous during the following weeks. This qualitative assessment was confirmed by computing differences between pre- and post-stroke FC across timepoints (Fig. S2A) and testing for significant changes in correlations using the network-based statistic(27,28) (Fig. 2D). Interestingly, a hypo-connected interhemispheric network emerged early and persisted up to 28 days after stroke. In contrast, the intra-hemispheric FC of the lesioned side showed a transient increase in the peri-infarct regions during the sub-acute phase followed by a gradual reduction in the chronic phase. One month after stroke, the hypo-correlated network expanded over the entire dorsal cortex, involving also distant regions like visual areas (Fig. 2D). Removing the correction for the hemodynamic signal did not significantly affect these findings (Fig. S2 B-C), in line with previous results from Nakai and colleagues.(28)

**Fig. 2.**
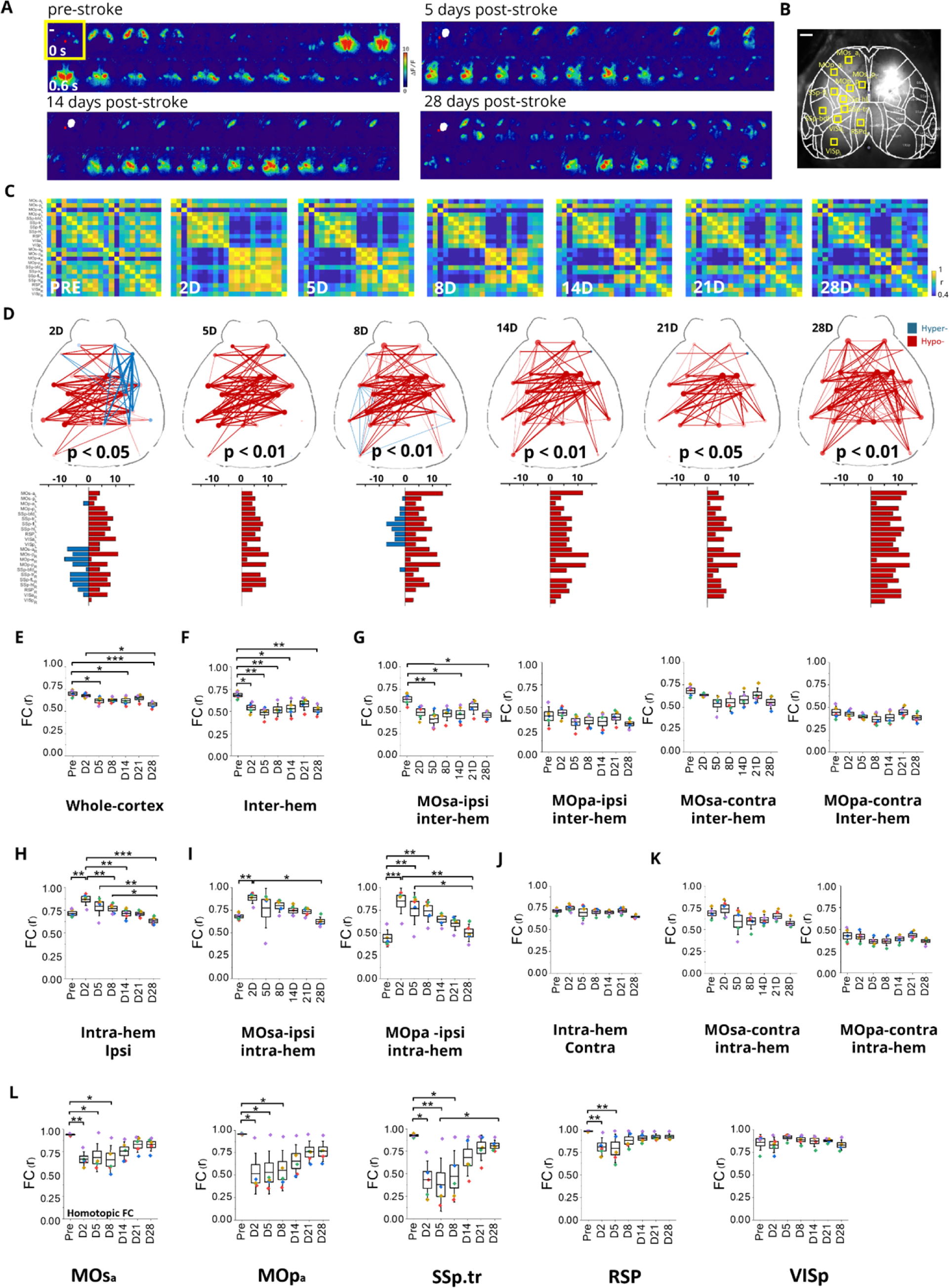
Enduring reduction of resting state FC of PV-IN after stroke. **A,** Representative image sequence of cortical PV activity before (upper left) and 5, 14 and 28 days after stroke. The white area locates the damaged region. The red dot indicates bregma (Scale bar, 1 mm). **B**, Wide-field calcium imaging field-of-view aligned with the surface of the Allen Institute Mouse Brain atlas. The white area on the left hemisphere locates the damaged region. Yellow squares represent cortical areas defined in both left (L) and right (R) hemispheres. Blue dots indicate bregma and lambda (Scale bar, 1 mm). **C**, Pairwise Pearson’s correlation coefficients of cortical activity were visualized as averaged correlation matrices pre-stroke and 2, 5, 8, 14, 21 or 28 days after injury. **D**, Network diagrams of statistically significant FC alterations after 2, 5, 8, 14, 21 or 28 days from injury. Blue and red lines denote significant hyper-correlation and hypo-correlation compared to pre-stroke values, respectively. The bar plots (bottom) indicate the number of significant FC alterations for each cortical area. **E**, Box chart illustrating FC averaged over the whole cortex. **F**, Box chart illustrating the averaged inter-hemispheric FC. **G**, Box charts showing inter-hemispheric FC of ipsilesional (left) and contralesional (right) secondary and primary motor cortices in the anterior regions (MOsa and MOpa respectively). **H**, Intra-hemispheric FC of ipsi-lesional areas. **I**, Box charts displaying intra-hemispheric FC of the secondary (left) and primary (right) ipsilesional motor cortices in the anterior region. **J**, Intra-hemispheric FC of contralesional areas. **K**, the box charts displaying intra-hemispheric FC of the secondary (left) and primary (right) contralesional motor cortices in the anterior region. **L**, Homotopic FC changes from pre-stroke to 28 days after injury (MOsa, anterior secondary motor cortex; MOsp, anterior primary motor cortex; SSp.tr, primary somatosensory cortex-trunk; RSP, dorsal part of the retrosplenial cortex; VISp, primary visual cortex). One Way ANOVA followed by Tukey test, * *P* < 0.05, ** *P* < 0.01, *** *P* < 0.001. Data are shown as mean ± SEM. Each color indicates a single subject, *n* = 5.

By quantifying the whole-cortex FC across time, we confirmed that the global reduction was not recovered 28 days after stroke (Fig. 2E). Then, the contribution of inter- and intra-hemispheric FC to the global decrease was evaluated separately. Interestingly, inter-hemispheric connectivity was dominated by a decrease in correlation strength (Fig. 2F). Notably, the anterior portion of the ipsilesional secondary motor cortex is the only motor area to display a significant FC decrease following stroke (Fig. 2G). Instead, intra-hemispheric FC shows a large but transient increase ipsilateral to the injury site (Fig. 2H), with a strong contribution from both the anterior portion of the peri-infarct primary and secondary motor cortices (Fig. 2I). Instead, there were no significant changes in the contralateral intra-hemispheric FC (Fig. 2J and 2K). Overall, these results imply that the decrease in global strength is mostly brought on by inter-hemispheric alterations. Therefore, we specifically dissected the contribution of homotopic desynchronisation: substantial decrease of homotopic connectivity was primarily affecting the peri-infarct and associative regions (M2, M1, SSp.tr, RSP) in the sub-acute phase, followed by a partial recovery starting a week from the damage (Fig. 2L). Our results suggest profound alterations in PV connectivity, which are not restricted to the injured site but extend throughout the entire dorsal cerebral cortex affecting primarily inter-hemispheric strength. Collectively, these alterations result in a consolidated hypo-connected network 28 days after stroke, indicating a failure of spontaneous recovery in the chronic phase.

### PV-IN in perilesional premotor cortex involved in motor execution can be engaged by optogenetic stimulation

Once established the significant impact of ischemic injury in CFA on PV-interneurons activity in perilesional tissue, we investigated whether the PV involved in voluntary movement circuitry could still be engaged by external stimulation. We induced either an ischemic or a sham lesion in CFA of B6;129P2-Pvalb tm1(cre)Arbr/J (PV-CRE) mice previously injected with a dflox.hChR2 AAV in RFA. Motor function was evaluated with Gridwalk test pre-lesion and 2 days after lesion in order to assess motor impairment in stroke mice (Fig. 3B). Subsequently, recordings from the spared perilesional premotor cortex were conducted 5 to 7 days after ischemic or sham lesion in awake, head-restrained mice. First, we used 200 ms blue-light pulses to identify putative PV-IN that exhibited increased firing rate lasting for the entire duration of the stimulation and putative neurons connected to the stimulated PV-interneurons, whose spontaneous discharge was inhibited during the stimulation. Secondly, we monitored the identified neurons during the execution of the forelimb retraction task on the M-Platform (Fig. 3A). Out of 35 PV-interneurons identified in 6 healthy animals, 19 correlated positively or negatively with movement onset. In 4 stroke animals, 17 out of 34 putative PV-interneurons were movement-related (Fig. 3C and 3D). PV-interneurons in stroke animals exhibited a significantly reduced firing rate during stimulation (Fig 3E). These findings demonstrated that, despite the profound impairment of PV-interneurons network and cell responsiveness resulting from the ischemic lesion, individual cells retain their reactivity, their involvement in forelimb movement execution and could be recruited by external stimulation.

**Fig. 3.**
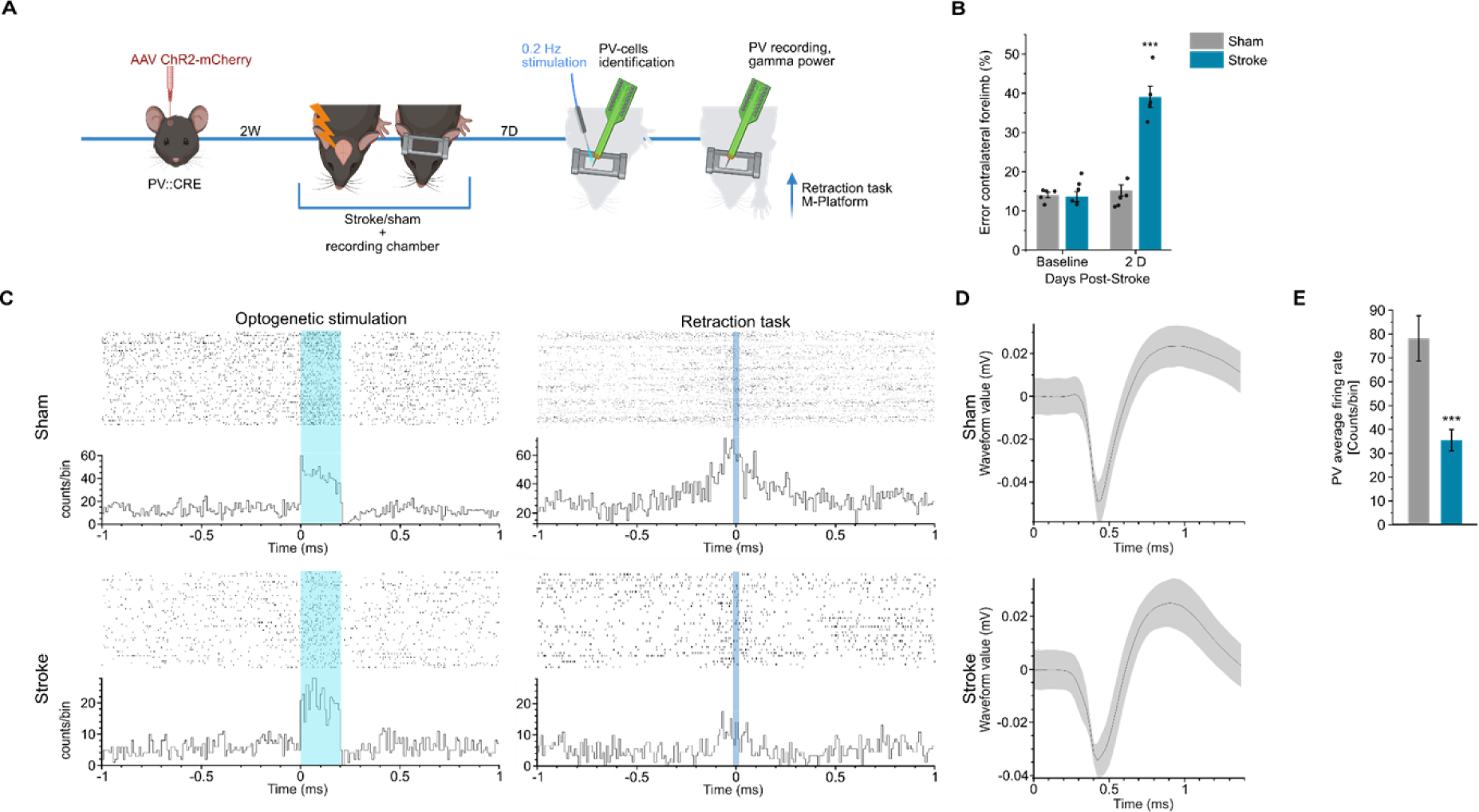
PV-IN in perilesional tissue respond to optogenetic stimulation and are involved in voluntary movement but show a decreased firing rate. Optogenetic stimulation and single-unit recordings were applied in order to identify PV-interneurons in the premotor cortex of healthy and ischemic animals and evaluate their discharge properties and their involvement in a voluntary movement. **A**, Schematic representation of the experimental procedure is reported. **B**, Motor function assessment in the Gridwalk test before and 2 days after stroke induction in sham (*n =* 5, gray bar plots) and stroke (*n =* 5, blue bar plots) animals, circles represent single animals (Two-Way ANOVA followed by Tukey test *** *P* < 0.001). **C**, Representative raster plots of identified PV-interneurons for sham and stroke animals. Upper panels display the raster plot of a PV-interneurons in response to optogenetic stimulation (left) and temporally aligned with movement onset (right) in a sham animal. Lower panels present the same rasters for a stroke animal. **D**, Average waveforms of recorded PV-IN for sham and stroke animals. **E**, Quantification of the average firing rate of the recorded PV-IN in response to optogenetic stimulation. A significant decrease in firing activity is evident in stroke animals (Two-tailed T-Test *** *P* < 0.001). Data are shown as mean ± SEM.

### Selective 40 Hz optogenetic PV-IN stimulation combined with robotic rehabilitation restores forelimb function

In order to determine the therapeutic potential of PV-interneurons-driven Gamma band stimulation, we used optogenetic stimulation in PV-CRE mice previously injected with a dflox.hChR2 AAV in RFA. We first demonstrated that 40 Hz optogenetic stimulation (3s pulses) increases Gamma band power in PV-CRE mice, but not in WT mice injected with dflox.hChR2 AAV, thus confirming the direct involvement of PV-interneurons activation in Gamma band generation (Fig. 4A). Subsequently, we investigated a combined neurorehabilitative approach where PV-CRE mice injected with dflox.hChR2 AAV were subjected to photothrombotic stroke in CFA. Mice were divided into three experimental groups: the Robot group received sham stimulation, the Robot 8 Hz group was treated with 8 Hz optogenetic stimulation in RFA during retraction task execution and the Robot 40 Hz group received 40 Hz stimulation. All groups received daily robotic rehabilitation on the M-Platform. Treatment efficacy was evaluated using the Gridwalk and Schallert Cylinder test, performed at baseline, 2 days after stroke (pre-treatment), once a week until 37 days post-stroke and after one week of follow-up without treatment. The experimental protocol schematic is shown in Fig. 4B. Initial deficits detected with both behavioral motor tests were consistent across groups (Fig. 4C and 4D). Robotic rehabilitation alone (grey bar plots) did not yield therapeutic benefits even after 5 weeks of treatment. Of note, combining 8Hz stimulation with Robotic rehabilitation (yellow bar plots) did not lead to additional motor improvement, indicating that PV activation at a different oscillatory pattern fails to restore motor function. Conversely, 40 Hz stimulation combined with robotic rehabilitation (blue bar plots) resulted in a highly therapeutic effect, evident both in the number of contralesional forelimb foot faults of the Gridwalk test and in the percentage of contralesional forelimb use of the Schallert Cylinder test. In both functional motor tests, the parameters reverted to baseline levels 37 days after injury. Importantly, the improvement remained stable during the follow-up assessment conducted one week after treatment conclusion. Post-mortem analyses showed no significant differences in PV-infected cells percentages among groups (Fig. 4E). In Fig. 4F, a magnification of the perilesional RFA of the Robot 40Hz group is reported as an example of double labeling of AAV-infected cells (in red) and immunohistochemically labeled PV-interneurons (in green).

**Fig. 4.**
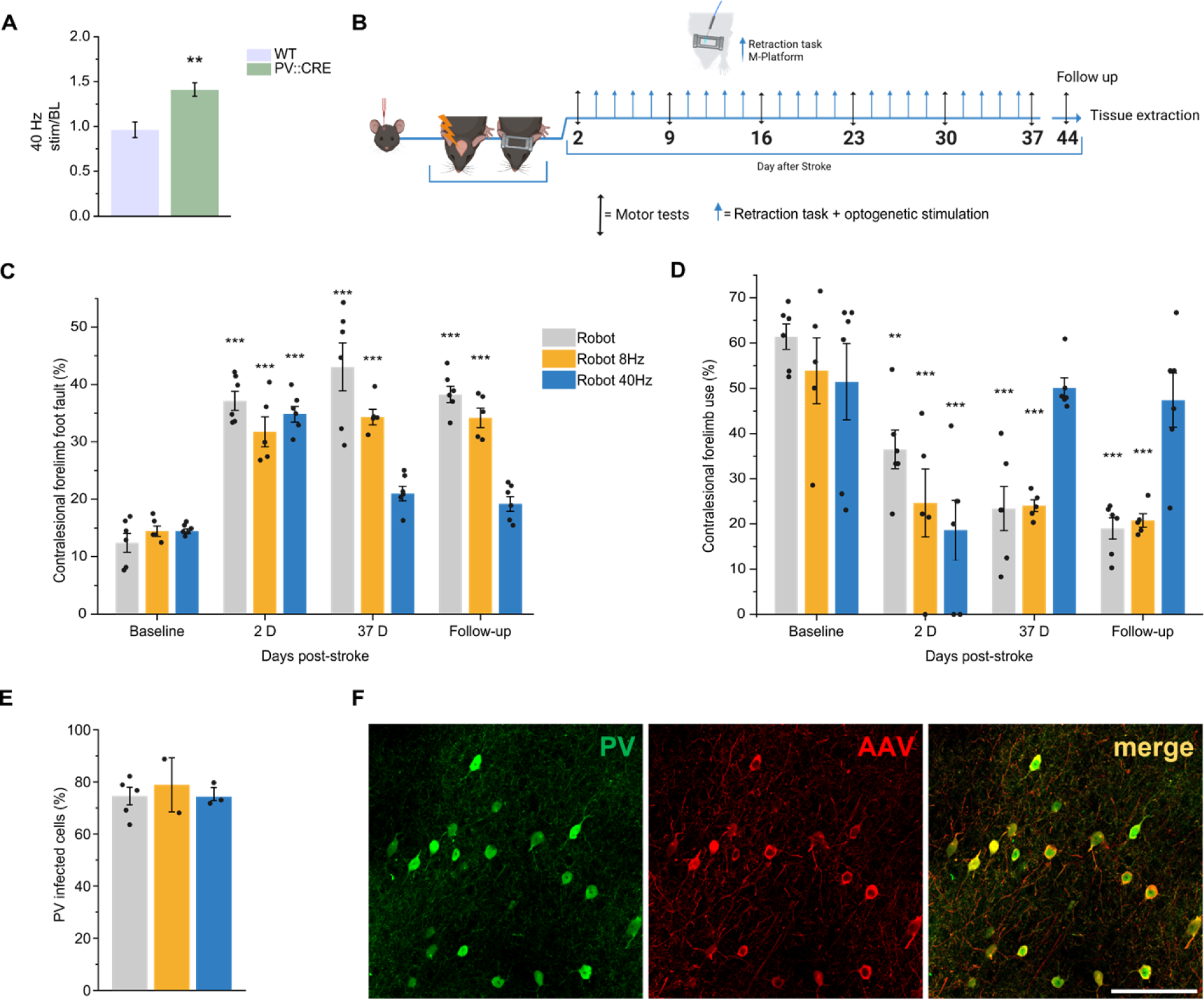
Robotic rehabilitation combined with optogenetic PV-interneurons stimulation at 40 but not 8 Hz improves motor function after stroke. **A**, Increase in Gamma band power in response to a 40 Hz optogenetic stimulation in PV-CRE mice (*n =* 5) but not in WT mice (*n =* 3) infected with dflox.hChR2 AAV (Student’s t-test, ** *P < 0.01*). **B**, Schematic of the experimental protocol. **C** and **D**, Forelimb motor function assessment in Gridwalk and Schallert cylinder test respectively. Robotic rehabilitation alone (grey bar plots, *n =* 6) is not able to restore forelimb motor function as expected. Similarly, 8 Hz stimulation (yellow bar plot, *n =* 5) fails to provide any improvement to motor rehabilitation. On the contrary, 40 Hz stimulation (blue bar plots, *n =* 6) significantly improved motor function (circles represent single animals). Two-Way RM ANOVA followed by Tukey test, * *P* < 0.05 ** *P* < 0.01, *** *P* < 0.001. Data are shown as mean ± SEM, circles represent single animals **E**, Quantification of PV-infected cells in the perilesional premotor cortex in the three experimental groups, revealing no significant differences among them (Robotic rehabilitation alone, *n =* 5; 8Hz stimulation, *n =* 2; 40 Hz stimulation, *n =* 3). One-way ANOVA followed by Dunnett’s test, *P* < 0.05. **F**, Representative image of PV-interneurons (in green), AAV infected cells (in red). Scale bar: 100 µm.

### Non-Invasive Gamma band stimulation in perilesional premotor cortex coupled with robotic rehabilitation improves forelimb motor function after stroke

We then evaluated whether the therapeutic effect obtained with specific optogenetic activation of PV-IN could be replicated using a more translational approach, the transcranial Alternating Current Stimulation (tACS). To this end, a photothrombotic stroke was induced in the CFA of a total number of 12 C57Bl6/J mice, which were subsequently divided into two experimental groups. All animals underwent 5 weeks of daily robotic rehabilitation. The Robot tACS group received 40 Hz tACS applied over the perilesional RFA, while the Robot group underwent sham stimulation, entailing electrode insertion and connection without actual current transmission (Fig. 5A). Motor function, assessed using Gridwalk and Schallert Cylinder tests, revealed significant deficits 2 days after the ischemic lesion induction, while no differences were observed between Robot tACS group and Robot group before lesion and 2 days post-lesion. Robotic rehabilitation alone did not lead to recovery. However, the Robot tACS group displayed significant improvements, returning to baseline in both tests and maintaining these gains during the follow-up period (Fig. 5B and 5C).

**Fig. 5.**
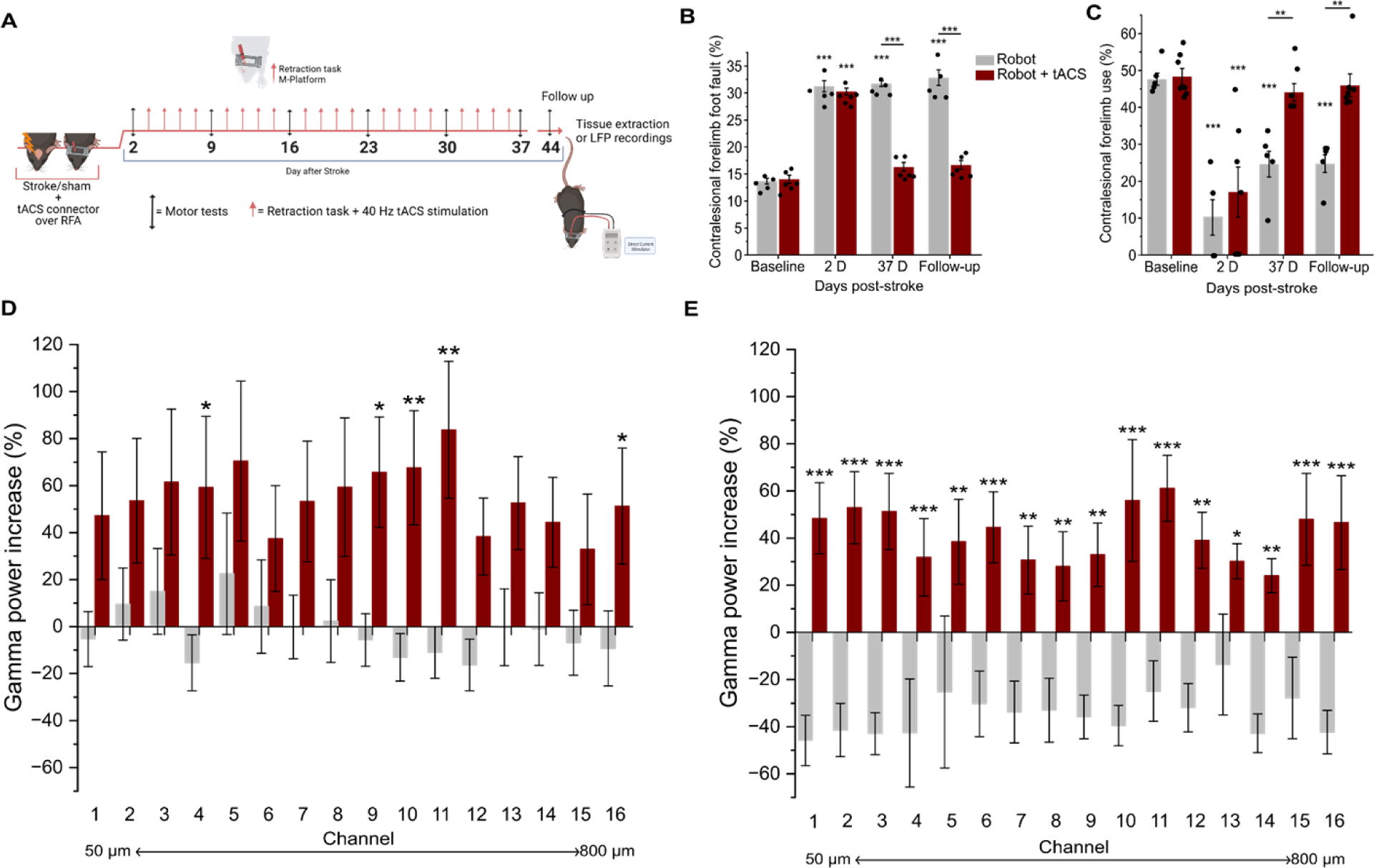
Forelimb motor improvements can be achieved by coupling robotic rehabilitation with non-invasive 40 Hz tACS. **A,** Schematic of the experimental protocol. **B** and **C**, Motor function assessment in mice treated with robotic rehabilitation alone (grey bar plots, *n =* 5) and coupled with non-invasive 40 Hz tACS (red bar plots, *n =* 6) on the perilesional premotor cortex evaluated with Gridwalk test and Schallert Cylinder test respectively. The use of 40 Hz stimulation succeeded in restoring forelimb motor function and maintained this improvement in a follow-up measure. **D** and **E,** Quantification of Gamma band power in pre-onset and post-onset windows respectively, across all 16 channels spanning the entire cortical layers at the follow-up time point (channel 1 ≃ 50 μm, channel 16 ≃ 800 μm). Grey bar plots *n =* 7, red bar plots *n =* 6. Two-Way RM ANOVA followed by Tukey test, * *P* < 0.05 ** *P* < 0.01, *** *P* < 0.001. Data are shown as mean ± SEM, circles represent single animals.

Next, we evaluated whether tACS treatment could induce a recovery in Gamma band oscillations in a new cohort of C57Bl6/J mice subjected to photothrombotic stroke in the CFA. Mice were divided into two experimental groups: the Robot-tACS group underwent 40 Hz tACS combined with robotic rehabilitation, while the Robot group received robotic rehabilitation alone. At the follow-up time point, the Robot-tACS group exhibited a significant increase in Gamma band power in both the pre-onset and post-onset windows (Fig. 5D and E), suggesting that the combined treatment approach, not only improves motor function, but also has a substantial impact on power within the Gamma frequency range. These findings underline the potential benefits of integrating robotic therapy with tACS in order to promote neural recovery and rehabilitation.

### Combined neurorehabilitation increases PV-IN connections and modulates GABAergic system

Following the assessment of motor function, brain tissues from both Robot and Robot-tACS not implanted groups were collected for immunohistochemical analysis in order to explore structural and biochemical impacts of the treatments. Focusing on the spared perilesional premotor cortex, we investigated changes in PV-interneurons synaptic connections by examining PV expression in “puncta rings” surrounding non-PV neuronal somas. The Robot-tACS group exhibited increased mean fluorescence compared to the Robot group, indicating enhanced PV-interneurons connectivity (Fig. 6A). Furthermore, we assessed whether the 40 Hz stimulation and the consequent increase in PV connections induces homeostatic changes in the GABAergic transporters in the perilesional cortex. We quantified the mean fluorescence of inhibitory terminals impinging on the soma of identified neurons. The expression of Vesicular GABA Transporter (VGAT) in the Robot-tACS group was significantly increased compared to Robot mice (Fig. 6B). These findings reveal that 40 Hz tACS stimulation significantly impacts the PV network and the GABAergic inhibitory system post-therapy.

**Fig. 6.**
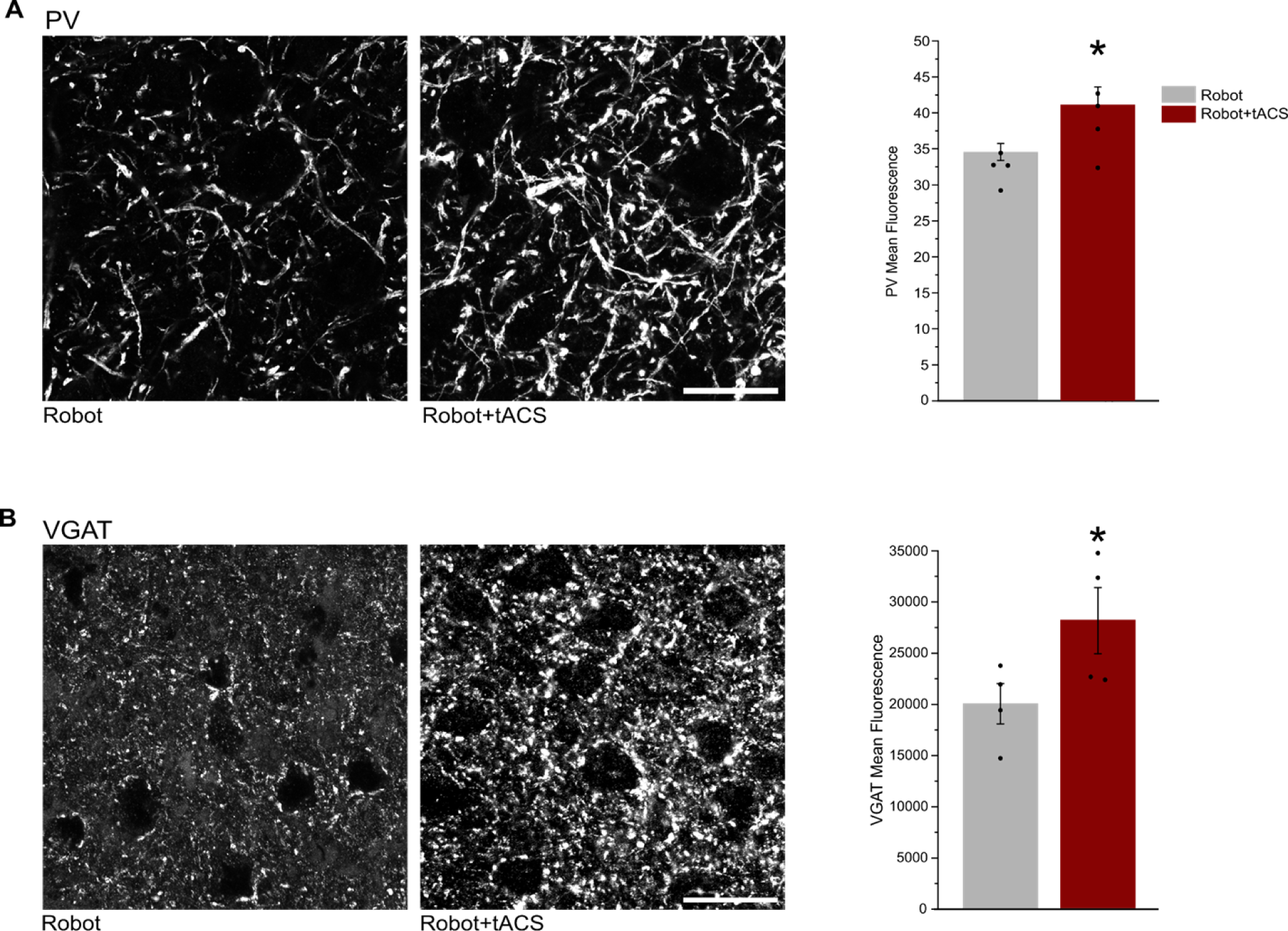
Increased PV-IN connections and modulation of GABAergic transporters after combined neurorehabilitation. Immunohistochemical characterization of the consequences of combined neurorehabilitation on the density of PV-interneurons connections and the expression of GABAergic transporters. **A**, Magnification of the perilesional premotor cortex immunostained with anti-PV for robot-only and robot+tACS groups together with the quantification of mean anti-PV fluorescence calculated in puncta-rings around cell bodies of non-PV positive neurons (unpaired Student’s t-test, * *P* < 0.05). Scale bar: 20 µm. **B**, Magnification of the perilesional premotor cortex immunostained with anti-VGAT antibody for robot-only and robot+tACS groups together with the quantification of mean anti-PV fluorescence calculated in puncta-rings around cell bodies of VGAT positive neurons (unpaired Student’s t-test, * *P* < 0.05). Scale bar: 20 µm. Data are shown as mean ± SEM.

## DISCUSSION

In this work, we delve into the role of PV-interneurons networks and movement-related Gamma modulation in post-stroke motor damage and rehabilitation in mice, offering novel and exciting insights that could significantly boost post-stroke treatment.

Our findings indicate significant alterations in Gamma modulation within the spared premotor cortex, both before and during voluntary movement. Consistently, following ischemic lesions, intra- and inter-hemispheric PV-interneurons connectivity becomes significantly impaired, worsening over time. Despite the reduced firing rate in perilesional tissue, survived PV-interneurons remain responsive to optogenetic stimulation, with their discharge consistently correlated with voluntary movement.These findings suggest that PV networks in perilesional surviving tissues may still be capable of triggering Gamma brain oscillatory activity tuned with movement control, if properly stimulated during physical rehabilitation. We then investigated a novel neurorehabilitative strategy, combining physical robotic therapy with 40 Hz optogenetic stimulation of PV-interneurons in spared perilesional tissue. With this protocol, we exploited the consistent action potential output of PV-interneurons with light pulses in the Gamma range (40 Hz)(20) and the compatibility of this stimulation of the ChR2 channel kinetics.(29–31) This approach specifically restored forelimb function in motor tests, while robotic rehabilitation alone or combined with 8 Hz stimulation, showed no improvement, confirming our previous findings.(1–3) Notably, coupling a 40 Hz clinically-relevant tACS stimulation with robotic rehabilitation also led to significant motor improvements, persisting even after one week of follow-up. Importantly, the treatment also restored movement-related Gamma band activity and subsequent post-mortem analyses, revealed increased PV-interneurons synaptic connections and altered GABAergic transporter expression, underscoring plastic functional and anatomical changes in the spared brain tissue.

Oscillatory brain activity in the Gamma range effectively engages brain areas in a cognitive task and has been recently associated with motor control and execution.(32,33)(34–36)(37) We recorded cortical activity in the perilesional forelimb premotor cortex during a forelimb retraction task and observed Gamma power modulation consistent with human studies,(34–36) underlining the premotor cortex’s role in movement planning and execution. Gamma power was positively modulated across all cortical layers during the pre-movement window, reflecting the recruitment of the premotor cortex in the task and computational processes related to movement preparation. During movement execution, the Gamma band remained positively modulated in the higher layers, while lower layers exhibited no modulation compared to the non-movement condition. This difference in Gamma power across layers could reflect distinct roles of higher and lower layers, with the former deputed to a computational engagement and the latter containing the larger pyramidal neurons conveying movement execution information to subcortical stations.(38,39) After stroke, Gamma band modulation in the motor cortex was significantly impaired, both before and during movement execution. Remarkably, no distinctions were observed between stroke and sham mice in the baseline window, emphasizing that frequency modulation deficits only emerge during active movement.

To correlate these findings with PV-interneurons connectivity alterations, we employed longitudinal Wide-Field imaging in PV-CRE mice infected with a viral vector expressing a floxed calcium indicator. Our results indicated significant impacts on PV activity and connectivity during the acute phase, primarily in peri-infarct cortical regions, with global hypoconnectivity persisting into the chronic phase of spontaneous recovery. Previous works using wide-field calcium imaging have underscored PV-interneurons’ crucial role in motor learning.(32,33) Despite their sparsity, inhibitory cells play a predominant role in many key brain functions, including neural network coordination(40) and memory formation.(41) Recently, a study focused on the large-scale connectivity of PV-interneurons demonstrated a loss of homotopic connectivity in stroke animals.(42–44) Here, we add a new piece to the puzzle and show that PV FC over the entire cortex is also compromised by a focal lesion, primarily on peri-infarct and homotopic regions. Stroke-induced de-differentiation of cortical activation may explain the transient increase in ipsilesional connectivity in the sub-acute phase. The reduced specificity in functional activation(45) might be advantageous in stroke subjects to activate brain regions nearby the lesion when executing functions that would normally require the damaged area. Therefore, hyperconnectivity could result from synchronous activation of the entire lesioned cortex. De-differentiation and synchronous activation of the entire cortex has already been shown to occur specifically in excitatory cortical neurons during the acute and chronic phases in spontaneously recovering stroke mice.(1) Collectively, these findings suggest that the excitatory/inhibitory balance is significantly disrupted at both local and distal levels, beginning in the acute phase and stabilizing in the chronic phase. In a previous study from our group, we demonstrated that post-stroke rehabilitation can alter the FC of excitatory neurons.(1) A recent study highlighted the cortex-wide properties of PV networks in a rat model of epilepsy,(46) and our research contributes further to this knowledge by addressing large-scale activity of PV-interneurons post-stroke. Given the crucial role of inhibitory neurons in other neurological conditions, including autism and Alzheimer disease,(47,48) we foresee that the same approach could be applied to study large scale alterations of inhibitory circuits in these and other neuropathologies.

Impairments in PV-interneurons connectivity could correlate with deficits in Gamma band modulation, as revealed by electrophysiology. Therefore, we investigated whether such dysfunctions could critically affect PV-interneurons responsiveness to external stimulation or voluntary movement. It has been demonstrated that PV-interneurons play an active role in voluntary movement execution through inhibition of pyramidal neurons,(49,50) and dysfunction in cortical PV networks has been reported in motor pathologies.(51–54) Combined with our wide-field imaging results, these findings suggest an early and long-lasting PV network dysfunction, potentially affecting their recruitment as targets for neurostimulation. Using optogenetic stimulation and single-unit recordings in PV-CRE head-fixed mice, we identified PV-interneuron activity during voluntary movement. Isomura and colleagues(49) identified representative fast-spiking interneurons as PV-IN correlated with lever pulling in mice. More recently, Giordano et al.*^57^* demonstrated that fast-spiking interneurons in the motor cortex and RFA display unique discharge properties, firing earlier and longer during movement compared to pyramidal neurons. Accordingly, we found many cells responding to optogenetic stimulation manifesting spiking activity aligned with movement onset in both control and stroke animals. This confirms active, movement-related PV-interneurons in the perilesional premotor cortex that can be recruited for movement-related Gamma band stimulation shortly after stroke. However, the ischemic lesion appears to compromise the physiological spiking activity of these cells, underlined by a decrease of firing rate identified in our recordings. This aligns with our previous findings of decreased numbers of PV-interneurons and their connections impinging on pyramidal neurons in the perilesional cortex of chronic stroke animals.(2,3,55,56) This underscores the importance of timing in interventions, confirming a critical window for cortical plasticity(57–59) that has to be exploited for rehabilitative purposes.

Considering these results and recognizing the importance of physical therapy in guiding plastic phenomena, we tested a combined neurorehabilitative approach involving Gamma band stimulation and PV-interneurons preservation in our mouse model. These methods have been validated both in clinical settings(60,61) and in animal models.(62) We focused on non-invasive neuromodulation techniques for their translational potential,(63) particularly tACS, which can entrain specific brain rhythms at rest or during a specific task, thus requiring minimal current to achieve desired effect.(64) Moreover, it is well-established that tACS is more effective when applied during tasks aligned with the target frequency of brain activity.(65,66) This makes tACS protocols fit well with the crucial necessity of physical rehabilitation for stroke patients. Robotic rehabilitation, in particular, can allow a consistent amount of work and a current acquisition of kinetic and kinematic data. Here, we implemented a protocol combining daily robotic rehabilitation with Gamma band stimulation through direct PV-IN activation using optogenetics. Initiating the protocol 5 days post-stroke, during the subacute phase, allowed us to exploit the plastic critical period and accommodate the physiological delay necessary for engaging in rehabilitative exercises, mandatory for human patients. This clinically relevant approach led to increased performances in forelimb functional tests due to Gamma frequency stimulation with sustained improvements even after a period without rehabilitative intervention. We also demonstrated the feasibility of achieving similar outcomes using a more translational approach with non-Invasive Gamma tACS over the perilesional premotor cortex, effective in increasing Gamma power during active movement in treated mice. The success of this protocol paves the way for validating and implementing a similar, well-tolerated and non-invasive and rehabilitation protocol in clinical practice.

We also explored the morphological and functional changes induced by combined rehabilitation within the perilesional premotor cortex. Our findings indicate that Gamma band stimulation significantly influences PV-IN connections, supporting the intertwined relationship between Gamma band oscillations and the excitation/inhibition balance(19,20) within interconnected networks of GABAergic interneurons and pyramidal cells in primary motor cortex.(15,17,67) Gamma band oscillations can induce homeostatic changes in the inhibitory system, as evidenced by immunohistochemical analysis revealing increased GABAergic transporter expression following treatment. This finding suggests enhanced cellular responsiveness to the strengthening of the parvalbumin circuit following neurorehabilitation. Increased GABAergic transmission might also mitigate further neuronal death during the acute phase. However, in the chronic phase, the role of GABA becomes more complex. Alterations in GABA release post-stroke can considerably alter the expression of GABA-A receptors, the main players in GABAergic transmission, leading to modification in both synaptic (phasic) and extrasynaptic (tonic) inhibition and to conflicting results in the literature. Recent studies, such as Clarkson et al.(68), have focused on manipulating tonic inhibition, showing that decreasing it after stroke improves motor performances in mice. These results were also confirmed in our laboratory, where modulating presynaptic GABA signaling in the first week after stroke in a mouse model improves long term motor functions.(55) However, literature reports conflicting effects when modulating phasic inhibition,(69) reporting an increase in phasic inhibition in perilesional areas during the critical window of cortical plasticity in a mouse model of stroke. By positively modulating this phasic inhibition with Zolpidem, an improvement in the motor performance of the animals was obtained. More recently, a selective increase of phasic inhibition has been demonstrated after a treatment with “continuous theta bursts simulation” in mice with an ischemic lesion and a corresponding functional improvement. These results suggest a positive role of increased phasic inhibition in the post-acute phase of ischemic injury but this topic is still debated.(70)

In summary, our study demonstrates the effectiveness of a clinically valuable novel neurorehabilitation approach combining physical therapy with movement-related Gamma band stimulation. This innovative strategy offers promising avenues for enhancing post-stroke recovery and deepens our understanding of the intricate dynamics of neural rehabilitation.

### Limitations of the study

This study raises many talking points and leaves many issues to be studied deeper and with different techniques. The use of animal models presents some disadvantages, due to differences in nervous system complexity and brain size, which can affect the precision and applicability of tACS treatment compared to human applications. Additionally, the precise molecular and electrophysiological mechanisms of tACS are not entirely understood and the effect observed at a molecular point of view requires further investigations. Moreover, the photothrombotic lesion method used in this study, although advantageous in terms of reproducibility and spatial specificity, does not perfectly mimic human ischemic pathology due to its lack of a penumbra and highly focal nature. These issues underscore the need for ongoing refinement of these methods and their adaptation for clinical use. Nevertheless, the high translatability of the results obtained with tACS, and the well-tolerated nature of these neurostimulation techniques, coupled with the rapidly expanding use of robotic devices in post-stroke rehabilitation, encourages prompt clinical trials to validate and potentially implement this integrated rehabilitative approach.

## MATERIAL AND METHODS

### Study Design

This study included 32 C57BL6/J and 37 B6;129P2-Pvalb-tm1(cre)Arbr/J (PV::Cre, Jackson Laboratories, JAX stock #017320) adult mice, aged 2-3 months. Animals were housed under standard conditions with a 12 h light/dark cycle and free access to food and water. Experimental procedures adhered to the ARRIVE guidelines and the European Communities Council Directive #86/609/EEC, and were approved by the Italian Ministry of Health (260/2016-PR, dated 11/03/2016). The required sample size was estimated based on previous results*^23^* and power calculations using G Power Software (v3.1.5). Animals were assigned randomly to groups.

### Ischemic Lesion

Cortical ischemic damage in the Caudal Forelimb Area (CFA) was induced using the photothrombosis method, as previously reported(3) and better described in Supplementary material. Following the photothrombotic procedure, animals underwent a head-restraining implantation surgery, consisting in a metal L shaped bar posted on the occipital bone using dental cement (Super Bond C&B, Sun Medical Company, Japan). For electrophysiology, a metal screw connected to a ground electrode was implanted in the occipital bone and a craniotomy was performed to expose the rostral forelimb area (RFA, 2.0 mm anterior and 1.25 mm lateral to Bregma(71)). The recording chamber was created by encircling the craniotomy hole with dental cement, then covered with agarose (1% in physiological solution) and silicon (Kwik-Cast Sealant, World Precision Instruments, USA) to protect the underlying tissues. For optogenetics, an optical fiber was positioned over the craniotomy. For tACS, a plastic tube was cemented over the RFA.

### Wide-field calcium imaging

#### Surgery

One week after AAV injection, mice were implanted with an intact skull preparation to allow free optical access to the cortex (modified from(2,72)). The skin and the periosteum were removed. Bregma was marked for stereotactic reference. A custom-made aluminum head-bar placed behind lambda was glued to the skull using transparent dental cement (Super Bond C&B – Sun Medical). The exposed cortex was then covered with the same cement. One week after the surgery, mice were habituated to head fixation under the wide-field microscope before the first imaging session. Calcium imaging was performed on PV-CRE mice the week before and 5 to 30 days after stroke.

#### Image processing and data analysis

Image stacks for each animal collected from different sessions were registered using custom-made software, by taking into account the bregma and λ position. An animal-specific field of view template was used to manually adjust the imaging field daily. To dissect the contribution of each cortical area, we registered the cortex to the surface of the Allen Institute Mouse Brain Atlas (www.brain-map.org) projected to our plane of imaging. For each block, image stacks were processed to obtain the estimates of ΔF/F0 (see Supplementary Materials).

A total of 22 ROIs were then selected (11 ROI for each hemisphere, 20×20 pixels). Correlation mapping was done for each subject by computing Pearson’s correlation coefficient between the average signals extracted from each ROI, with that of each other ROI. The single-subject correlation maps were then transformed using Fisher’s r-to-z transform and then averaged across all animals. Averaged maps were re-transformed to correlation values (r-scores). For each mouse, r(pre-stroke)-r(x-day post-stroke) was calculated and averaged across mice in order to visualize matrices of difference between the pre-stroke condition and all the time points after-stroke.

### Forelimb retraction task on the M-Platform

The M-Platform(1,26) is described in detail in Supplementary Materials. Daily training sessions were conducted for both electrophysiological recordings and rehabilitation experiments. Each session involved 15 forelimb retractions by each mouse, combining passive (device-extended by 10 mm) and active (animal-retracted) movements. Overcoming a force threshold earned the mice a liquid reward for each successful task completion. Mice typically mastered this task, improving performance within 2-3 days.(26) Rehabilitation began 5 days post-lesion and continued until day 37, occurring for 4 consecutive days a week. Adjustments to task friction were made according to each animal’s functional deficit. Details on coupled sham or neuromodulatory treatments are provided below.

### Viral Injection

For Parvalbumin Interneurons (PV-IN) identification and optogenetic Gamma band stimulation, PV::Cre transgenic mice received a stereotactic injection of the AAV vector (AAV1.EF1.dflox.hChR2(H134R)-mCherry.WPRE.hGH (Addgene, USA) to induce a Cre-dependent expression of Channelrhodopsin-2 (ChR2). The procedure is described in detail in Supplementary Material.

### Optogenetic stimulation

Optogenetic stimulation was employed for identifying PV-IN during electrophysiological recordings and as a neuromodulatory treatment for rehabilitation. The stimulation setup included a PlexBright Optogenetic Stimulation System with an LD-1 Single Channel LED Driver and a 456 nm LED Module (PlexonInc, USA), connected to a 200 µm Core 0.39 NA optic fiber (ThorlabsInc, USA). The maximum emission power of the optic fiber was assessed before each experiment with the PlexBright Light Measurement Kit. Control of stimulation parameters was managed via custom software in LabWindows/CVI, interfacing through a NI USB-6212BNC DAQ board (National Instruments, USA). For optogenetic identification of PV-IN, single 200 ms light pulses at 0.2 Hz were given at increasing light power. For optogenetic induction of Gamma band intrease, 40 Hz or 8 Hz, 1 ms light pulses were administered throughout the task—from the passive to retraction phases on the M-Platform. Task duration varied between 5 and 8 minutes, depending on each animal’s ability to complete a retraction.

### Electrophysiological recording

In C57Bl6/J and PV-ChR2 mice, Gamma power modulation and PV-IN discharge properties in the RFA were assessed using 16-channel linear probes (NeuroNexus, USA). Mice were head-fixed on the M-Platform, with the left forepaw linked to a load cell, and electrodes were stereotactically inserted in the right hemisphere’s RFA at a depth of 850 µm.

Neural signals were acquired and amplified using DigiAmp (Plexon, USA), with ground in the cerebellum. Recordings were made during resting and retraction task. Offline analysis, performed with custom algorithms in NeuroExplorer (Plexon, USA) and Matlab, involved computing power spectral density (PSD) and synchronizing neural signals with the force signal from the platform. Isolated force peaks (force peaks with no movement within 2.5 seconds before and after the movement onset) were selected. Peri-events spectrogram analysis was used to calculate the power within the 31-49 Hz frequency band in 0.5-seconds intervals at 3 different time windows relative to the onset of the force peak : Baseline (−2 to −1.5 seconds), Pre-onset (−0.5 to 0 seconds), and Post-onset (0 to 0.5). Ratios of Pre-onset / Baseline and Post-onset / Baseline were computed to evaluate Gamma band power variations in CFA during movement with respect to the No-Movement state. Gamma power variations were quantified as percentages of the total PSD, excluding the force peaks and the time windows immediately before and after the movement onset.

Optogenetic identification of PV-IN during resting states involved light pulses administered near the recording electrode. After the optogenetic stimulation protocol, neural activity was recorded during the retraction task without moving the recording electrode. To identify putative PV fast-spiking interneurons, whose firing rate increased selectively during optogenetic stimulation. Spike Sorting analysis for single-unit was performed offline using Offline Sorter Software (Plexon, USA) and was used to refine the principal component analysis (PCA) to exclude potential spikes originating from neighboring neurons. Once identified, putative PV-IN peri-event rate histogram was referred to the movement onset to verify firing rate alignment with voluntary movement. Recordings spanned 3 days to enhance identification of PV neurons involved in motor tasks.

### Behavioral motor tests

Functional assessment of forelimb motor performances was used to evaluate the impact of the ischemic damage and the applied neurorehabilitative protocols. Mice performances were assessed in baseline condition and then once a week at days 2, 9, 16, 23, 30, 37 and 44 post-lesion, as reported in Lai et al.(73). Two behavioral tests have been used, Gridwalk and Schallert Cylinder test, that are fully described in Supplementary Materials.

### Transcranial Alternating Current Stimulation (tACS)

tACS was administered using an AnimaltES Model 2101 (Soterix Medical Inc., USA) at a frequency of 40 Hz. The cathode was positioned in a saline-filled tube cemented to the skull, and the anode was placed under the abdomen against a saline-soaked sponge. Stimulation began 5 minutes prior to task onset, with current gradually increasing to a maximum of 0.2 mA, remaining below the movement threshold for mice. The stimulation continued for the task duration (5 to 8 minutes), with current ramp-down starting 10 minutes post-task initiation. Sham stimulation involved electrode placement without current flow. No signs of discomfort or freezing were observed in the animals during tACS.

### Histology

For immunostaining, brain slices were incubated in a blocking solution for 1 hour at room temperature (10% donkey serum; 0.3% Triton X-100 in PBS), treated with primary antibodies, prepared at the proper concentration in 1% donkey serum and 0.2 % Triton X-100 in PBS overnight at 4°C. Following 3 washes in PBS, the sections were incubated for 2 hours at room temperature with the specific secondary antibodies. A detailed description of this procedure is available in Supplementary Materials.

### Statistical considerations

Statistical analyses were conducted on raw data using SigmaPlot 11.0 (Systat Software Inc, USA), Matlab (R2019a), and OriginLab (2018), with a significance threshold set at alpha = 0.05. For behavioral tests (Gridwalk and Schallert Cylinder), Two-Way Repeated Measures ANOVA with Tukey post-hoc tests were applied. Group comparisons utilized One- or Two-Way ANOVA, depending on the data structure, followed by either Dunnett’s or Tukey’s post-hoc tests. T-tests were used for immunohistochemical analyses and firing rate comparisons. Group-level ROI-based functional connectivity (FC) differences pre- and post-stroke were analyzed using one-way repeated measure ANOVA with Tukey correction. Network Based Statistic (NBS) Toolbox in MATLAB assessed functional network connectivity.(27,28) Significance was determined at p < 0.05, and errors are expressed as Standard Error of Means (SEM), with significance markers (* P < 0.05, ** P < 0.01, *** P < 0.001).

## Data visualization

Data visualization was performed using OriginPro, while figure editing was performed with Inkscape 1.3.2. Cartoons were created with BioRender.com.

## Acknowledgments

We thank Francesca Biondi (CNR Pisa) for the excellent animal care and Elena Novelli with her technical support for the imaging section.

## Funding

Regione Toscana, PERSONA project, bando Salute 2018 (MC, SM)

H2020 EXCELLENT SCIENCE-European Research Council (ERC) under grant agreement 692943 (MC)

Regione Toscana, NIMBLE project (20RSVP), bando Salute 2018 (ALAM)

THE Tuscany Health Ecosystem ECS_00000017 MUR_ PNRR (ALAM)

Grant Fondazione Baroni 2021 (CS, MC)

## Author contributions

Conceptualization: CS, MC

Methodology: ALAM, CS, MC

Investigation: LV, FM, ALAM, EM, MP, AMa, AMi, EC, CS

Visualization: LV, FM, ALAM, EM, MP, SM, CS

Funding acquisition: SM, ALAM, MC

Project administration: CS, ALAM, SM, MC

Supervision: CS, ALAM, MC

Writing – original draft: LV, EM, ALAM, CS

Writing – review & editing: LV, FM AMi, ALAM, EM, MP, CS

## Data and materials availability

All source data are accessible in a public database.

